# Rewiring of the host cell metabolome and lipidome during lytic gammaherpesvirus infection is essential for infectious virus production

**DOI:** 10.1101/2023.01.30.526357

**Authors:** Sarah A. Clark, Angie Vazquez, Kelsey Furiya, Madeleine K. Splattstoesser, Abdullah K. Bashmail, Haleigh Schwartz, Makaiya Russell, Shun-Je Bhark, Osvaldo K. Moreno, Morgan McGovern, Eric R. Owsley, Timothy A. Nelson, Erica Sanchez, Tracie Delgado

## Abstract

Oncogenic virus infections are estimated to cause ∼15% of all cancers. Two prevalent human oncogenic viruses are members of the gammaherpesvirus family: Epstein Barr Virus (EBV) and Kaposi’s Sarcoma Herpesvirus (KSHV). We use murine herpesvirus 68 (MHV-68), which shares significant homology with KSHV and EBV, as a model system to study gammaherpesvirus lytic replication. Viruses implement distinct metabolic programs to support their life cycle, such as increasing the supply of lipids, amino acids, and nucleotide materials necessary to replicate. Our data define the global changes in the host cell metabolome and lipidome during gammaherpesvirus lytic replication. Our metabolomics analysis found that MHV-68 lytic infection induces glycolysis, glutaminolysis, lipid metabolism, and nucleotide metabolism. We additionally observed an increase in glutamine consumption and glutamine dehydrogenase protein expression. While both glucose and glutamine starvation of host cells decreased viral titers, glutamine starvation led to a greater loss in virion production. Our lipidomics analysis revealed a peak in triacylglycerides early during infection and an increase in free fatty acids and diacylglyceride later in the viral life cycle. Furthermore, we observed an increase in the protein expression of multiple lipogenic enzymes during infection. Interestingly, pharmacological inhibitors of glycolysis or lipogenesis resulted in decreased infectious virus production. Taken together, these results illustrate the global alterations in host cell metabolism during lytic gammaherpesvirus infection, establish essential pathways for viral production, and recommend targeted mechanisms to block viral spread and treat viral induced tumors.

**IMPORTANCE:** Viruses are intracellular parasites which lack their own metabolism, so they must hijack host cell metabolic machinery in order to increase the production of energy, proteins, fats, and genetic material necessary to replicate. Using murine herpesvirus 68 (MHV-68) as a model system to understand how similar human gammaherpesviruses cause cancer, we profiled the metabolic changes that occur during lytic MHV-68 infection and replication. We found MHV-68 infection of host cells increases glucose, glutamine, lipid, and nucleotide metabolic pathways. We also showed inhibition or starvation of glucose, glutamine or lipid metabolic pathways results in an inhibition of virus production. Ultimately, targeting changes in host cell metabolism due to viral infection can be used to treat gammaherpesvirus induced cancers and infections in humans.

## INTRODUCTION

The oncogenic viruses Epstein Barr Virus (EBV) and Kaposi’s Sarcoma Herpesvirus (KSHV) are both members of the human gammaherpesvirus family. Over 90% of the human population is infected with EBV, with an estimated 137,900 – 208,700 cancer related deaths in 2020 (1). Additionally, KSHV causes Kaposi’s Sarcoma (KS) in 1 out of every 200 transplant patients in the US (2). Lytic replication studies in both KSHV and EBV cell culture systems are challenging because both viruses must be reactivated from latently infected cells to induce lytic replication and they both cannot form plaques to quantify viral production (3, 4). Studying the induction of the lytic cycle during gammaherpesvirus infection is important as it is the key to spreading virus in the host, can seed new tumors, and has implications in cancer maintenance (5).

Murine Herpesvirus 68 (MHV-68), a natural pathogen of wild rodents, is a mouse gammaherpesvirus which shares ∼80% of its opening reading frames (ORFs) with EBV and KSHV (6). MHV-68 provides a genetically tractable model system that naturally undergoes both lytic replication and latent infection in various *in vitro* cell lines, while also forming plaques for quantitation of virus production. Overall, MHV-68 infection is an ideal model system to study gammaherpesvirus lytic replication. Ultimately, MHV-68 can be used in animal models to elucidate how gammaherpesviruses influence oncogenesis within the host.

Oncogenic viral infection modifies numerous host cell mechanisms, including metabolism. Viruses are intracellular parasites which lack their own natural metabolism and therefore must commandeer the host cell metabolic machinery to generate both energy in the form of ATP and other critical biosynthetic building blocks (7). Increasing energy and central carbon metabolism to generate more amino acids, lipids, and nucleotides is essential for viral replication and the formation of new virions. Previous work by our group and others have identified major changes in central carbon metabolism during viral infection. For example, RNA viruses, such as Dengue virus, Influenza virus, Zika virus, and SARS-CoV-2, and DNA viruses including Herpes simplex virus-1 (HSV-1), Human Cytomegalovirus (HCMV), Vaccinia Virus (VAVC), Adenovirus, and KSHV all induce significant changes in host cell glycolysis, glutamine metabolism, and fatty acid metabolism, revealing similarities and differences in modulating metabolism during infection (8–20). We previously showed latent KSHV infection of endothelial cells induces glucose, glutamine, and lipid metabolism, and that the induction of these metabolic pathways is necessary for the survival of latently infected cells (17–19). Studies of latent EBV infection have shown that methionine metabolism is key for maintaining latency (21). While critical alterations of metabolism during latent infection have been explored, metabolic changes during lytic infection with gammaherpesvirus are less well known (19, 22–25). A previous study of lytic KSHV infection showed that the induction of glycolysis, glutaminolysis, and fatty acid synthesis is required for viral replication, however no global analysis of metabolic alterations were reported (20). While we know these metabolic pathways are essential for maximal virus production, determining the viral mechanisms are important to identify potential therapeutic targets. Using MHV-68 as a strong genetically tractable system, it is possible to determine the cellular and viral mechanisms of metabolic modulation during gammaherpesvirus infections.

A few studies have begun to examine aspects of altered host metabolism during MHV-68 gammaherpesvirus infection. MHV-68 infected mice fed a high fat diet revealed MHV-68 infection promotes fatty liver and elevated triglyceride levels (26). Another study showed host cell cholesterol synthesis supports viral replication of MHV-68 in primary macrophages, and inhibiting cholesterol synthesis reduced virus production (27). A follow-up study in primary macrophages revealed that the regulation of cholesterol and fatty acid synthesis is in part due to the elevated expression of specific transcription factors called Liver X Receptors (LXRs) (28). A recent study further explored the role of hypoxia-inducible factor 1 alpha (HIF1ɑ) during MHV-68 infection, showing that lytic infection induces HIF1ɑ protein expression, a global regulator of cell metabolism, and blocking HIF1ɑ regulation reduces virus production and reactivation in infected mouse cells (29). Overall, these previous studies have highlighted individual metabolic pathways implicated in gammaherpesvirus virion production and opened the doors for further exploration of host cell metabolism changes during gammaherpesvirus lytic replication.

In this study, we define the global changes in the host cell metabolome and lipidome during gammaherpesvirus lytic replication and report that MHV-68 lytic infection induces glycolysis, glutaminolysis, lipid metabolism, and nucleotide metabolism. We also demonstrate that pharmacological inhibition of glycolysis or lipogenesis significantly reduce maximal MHV-68 viral production. Furthermore, we show that glutamine metabolism, rather than glucose metabolism, appears to be a more essential pathway for virion production. Understanding how the gammaherpesvirus lytic cycle plays a role in altering global host cell metabolism can provide us with new therapeutic mechanisms to block gammaherpesvirus lytic replication and virus spread.

## RESULTS

### Metabolomics analysis reveals the importance of nucleotide metabolism during gammaherpesvirus lytic replication

Metabolomics is a useful tool to determine global host cell changes during viral infections (8, 14, 17, 30–32). To profile the host cell metabolome during lytic MHV-68 infection, we harvested mock or MHV-68 infected NIH3T3 cells (MOI = 3) for targeted aqueous metabolomics analysis at 4, 8, 12, 24 and 36 hours post infection (hpi) in three independent experiments. Cells were harvested and rinsed in -80°C cold methanol and sent for targeted liquid chromatography - mass spectrometry (LC-MS) profiling at the University of Washington Northwest Metabolomics Research Center (NW-MRC). All metabolite concentrations were normalized by whole cell protein concentration. Fold change was calculated as a ratio of metabolite abundance in MHV-68 infected cells compared to mock infected cells.

A total of 176 metabolites were detected for all samples in biological triplicate at all time points; 24 lipid species, 69 amino acid species, 25 carbohydrate species, 36 nucleotide species, and 22 other species (Fig. 1 and Table S1). Nucleotide metabolism species contained the highest abundance of elevated metabolites throughout the virus life cycle compared to the other metabolic categories. Previous literature indicates MHV-68 infected cells at 24 hpi contain peak amounts of viral DNA and expression of genes encoding structural proteins and DNA packaging proteins (33, 34). Therefore, we analyzed the top statistically significant metabolites at 24 hpi (p<0.05 and at least 1.5 fold change). Volcano plot analysis depicting log 2 fold change (FC) and -log10 p-value (p) in 24 hpi samples (t-test, unpaired) highlighted 12 significantly elevated (red) and 3 significantly decreased (blue) metabolite species (Fig. 2A). Of the 12 elevated metabolites, 7 were nucleotide species and 2 others were nucleotide sugar species. Thymidine monophosphate (DTMP) was the most statistically significant elevated aqueous metabolite in our metabolomics screen, reaching 16-fold change over mock infected cells by 24 hpi (Fig. 2A and B, and Table S1).

**Figure 1:**
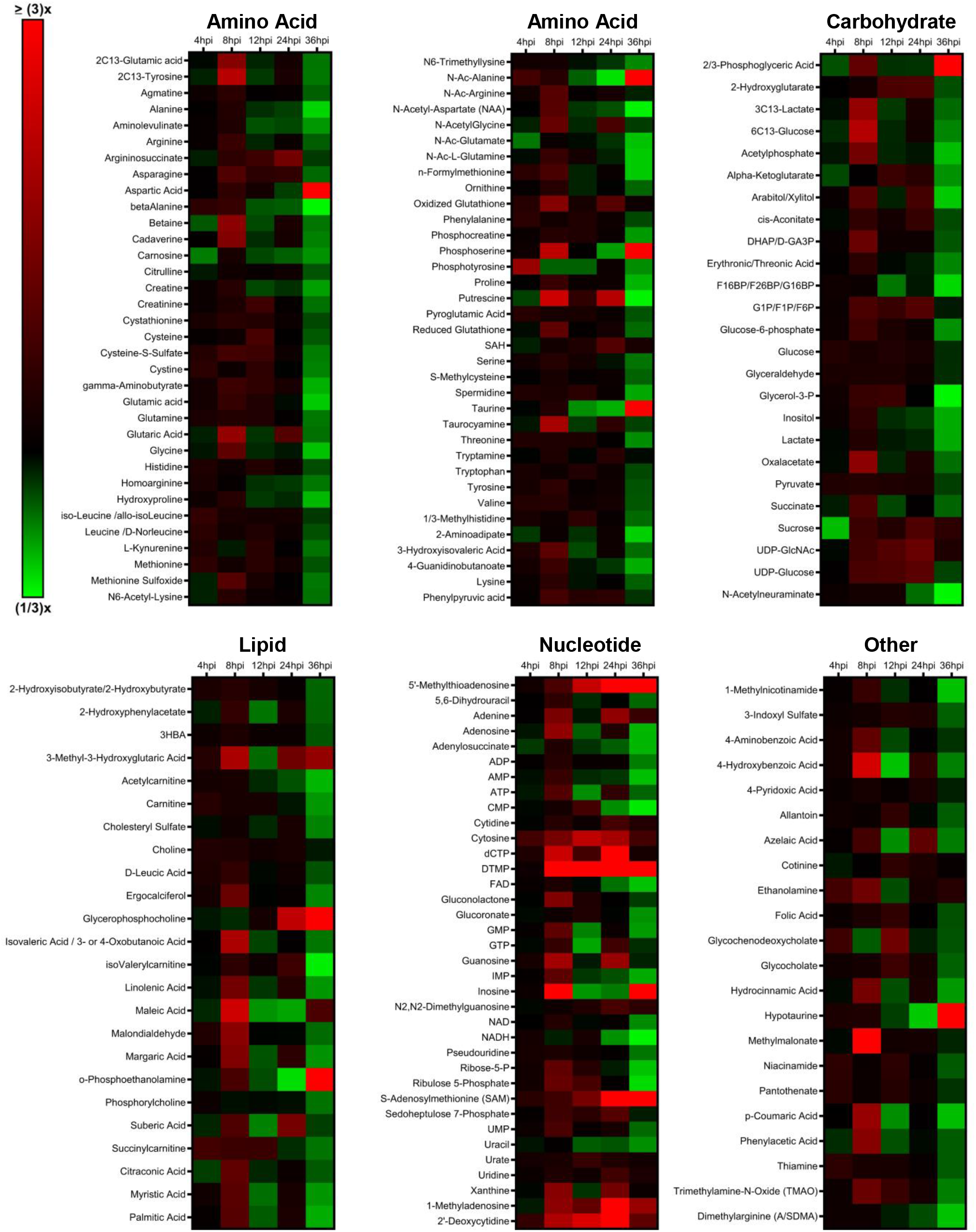
Targeted aqueous metabolomics of mock vs MHV-68 infected cells. Mock (control) and MHV-68 infected NIH3T3 cells were harvested at 4, 8, 12, 24, and 36 hpi in three independent experiments. Relative fold change time course depicted, where red shows an increase and green shows a decrease in metabolite concentration in MHV-68 infected cells compared to mock infected cells. Metabolites are organized according to the major metabolic categories (carbohydrate, lipid, nucleotide, and amino acid) or other.

**Figure 2:**
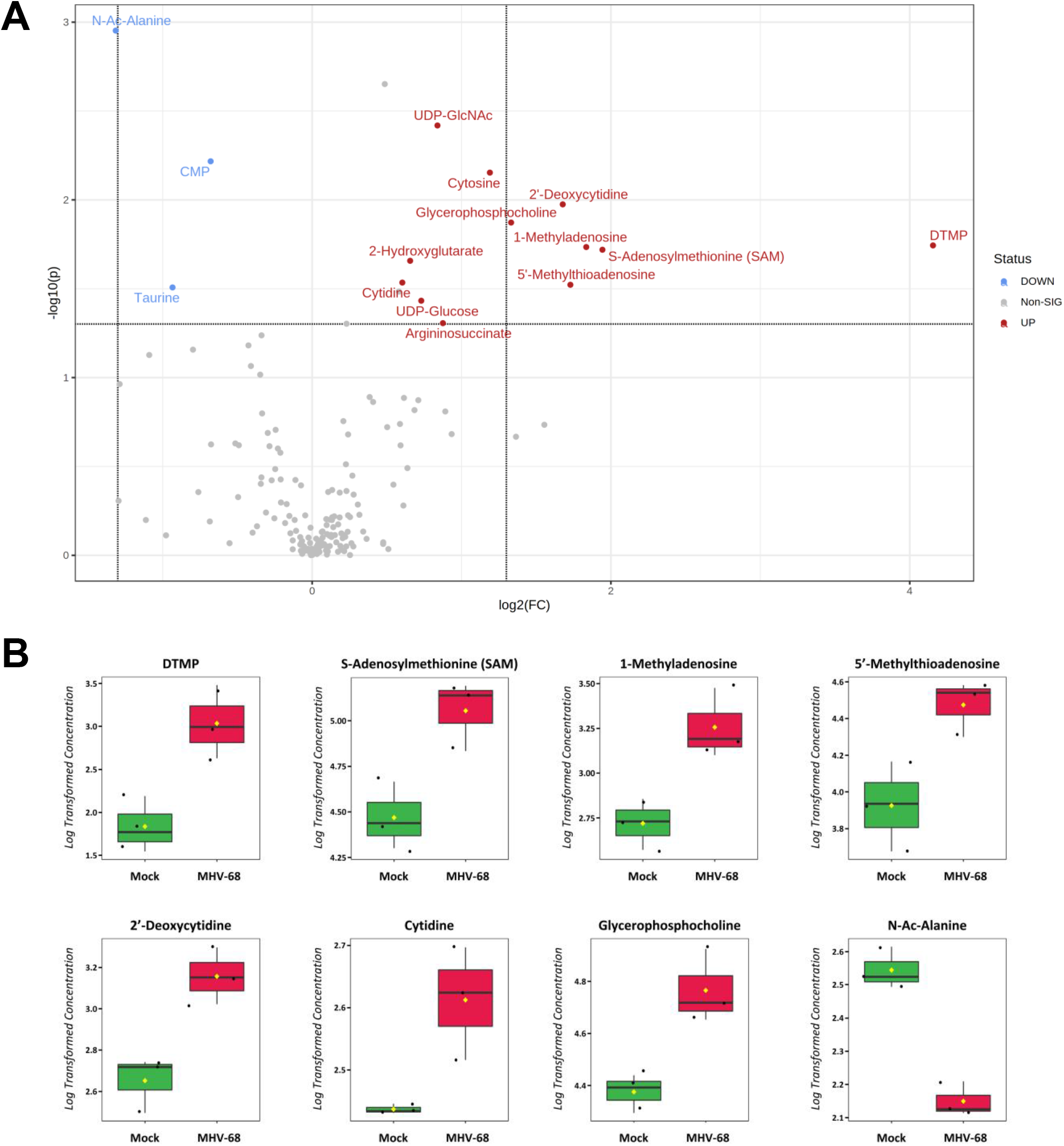
Aqueous metabolic profiling of mock vs MHV-68 infected cells at 24hpi. A) Volcano plot depicting log 2 fold change (FC) and -log10 p-value (p) in 24 hpi samples (unpaired). Statistical threshold set to p < 0.05 and at least 1.5 fold change in MHV-68 infected cells compared to mock infected cells using the student t-test. Red metabolites are significantly up and blue metabolites are significantly down. B) Box plots of the top 8 statistically significant metabolites (t-test, p<0.05, and at least 2 fold change). Black dots represent concentrations for all samples (biological triplicates).The notch indicates the median and yellow diamond indicates the mean concentration.

Additionally, there was elevation in other nucleotide species such as the nucleotide cytosine, two pyrimidine nucleosides, cytidine and 2’-deoxycytidine, and the purine nucleoside 1-Methyladenosine (Fig. 2A and B, and Table S1). S-adenosylmethionine (SAM) and its downstream metabolite 5’-methylthioadenosine (MTA), which belong to the same class of organic compounds (5’-deoxy-5’-thionucleosides), are linked to both adenine nucleotide and polyamine metabolism (35, 36) and were both elevated in infected cells (Fig. 2A and B, and Table S1). Uridine diphosphate glucose (UDP-glucose) and uridine diphosphate-N-acetylglucosamine (UDP-GlcNAc), both pyrimidine nucleotide sugars, were elevated in infected cells (Fig. 2A and Table S1). UDP-glucose is a key intermediate in carbohydrate metabolism and a precursor to glycogen, sucrose, lipopolysaccharides, and glycosphingolipids (35). UDP-GlcNAc is a fructose-6-phosphate and glutamine derivative, which is used as a substrate by glycosyltransferases to link N-acetylglucosamine to glycosaminoglycans, proteoglycans, and glycolipids (37, 38). Other elevated metabolite species in MHV-68 infected cells included 2-hydroxyglutarate (carbohydrate metabolism), glycerophosphocholine (lipid metabolism), and argininosuccinate (amino acid metabolism) (Fig. 2A and B, and Table S1). Our analysis also yielded only three significantly decreased metabolite species in MHV-68 infected cells compared to mock infected cells, including N-Ac-Alanine (amino acid metabolism), CMP (nucleotide metabolism) and taurine (amino acid metabolism) (Fig. 2A and B, and Table S1). Taken together, these data indicate that MHV-68 infected cells increase nucleotide metabolism during lytic replication at 24 hpi.

### MHV-68 lytic infection increases glucose metabolism

To determine if glucose metabolism is modulated during the MHV-68 lytic life cycle, we analyzed our aqueous metabolomics data for the abundance of metabolites in the glycolysis metabolic pathway and the TCA cycle. Fold change was calculated as a ratio of metabolite abundance in MHV-68 infected cells compared to mock infected cells. Our metabolomics analysis revealed MHV-68 infection increased all detected glycolytic intermediates at 8 hpi, an early part of the viral life cycle (Fig. 3A). Most TCA cycle metabolic intermediates also increased in abundance at 8 hpi, with oxaloacetate displaying the highest fold change in abundance out of all the TCA cycle metabolites (Fig. 3B).

**Figure 3:**
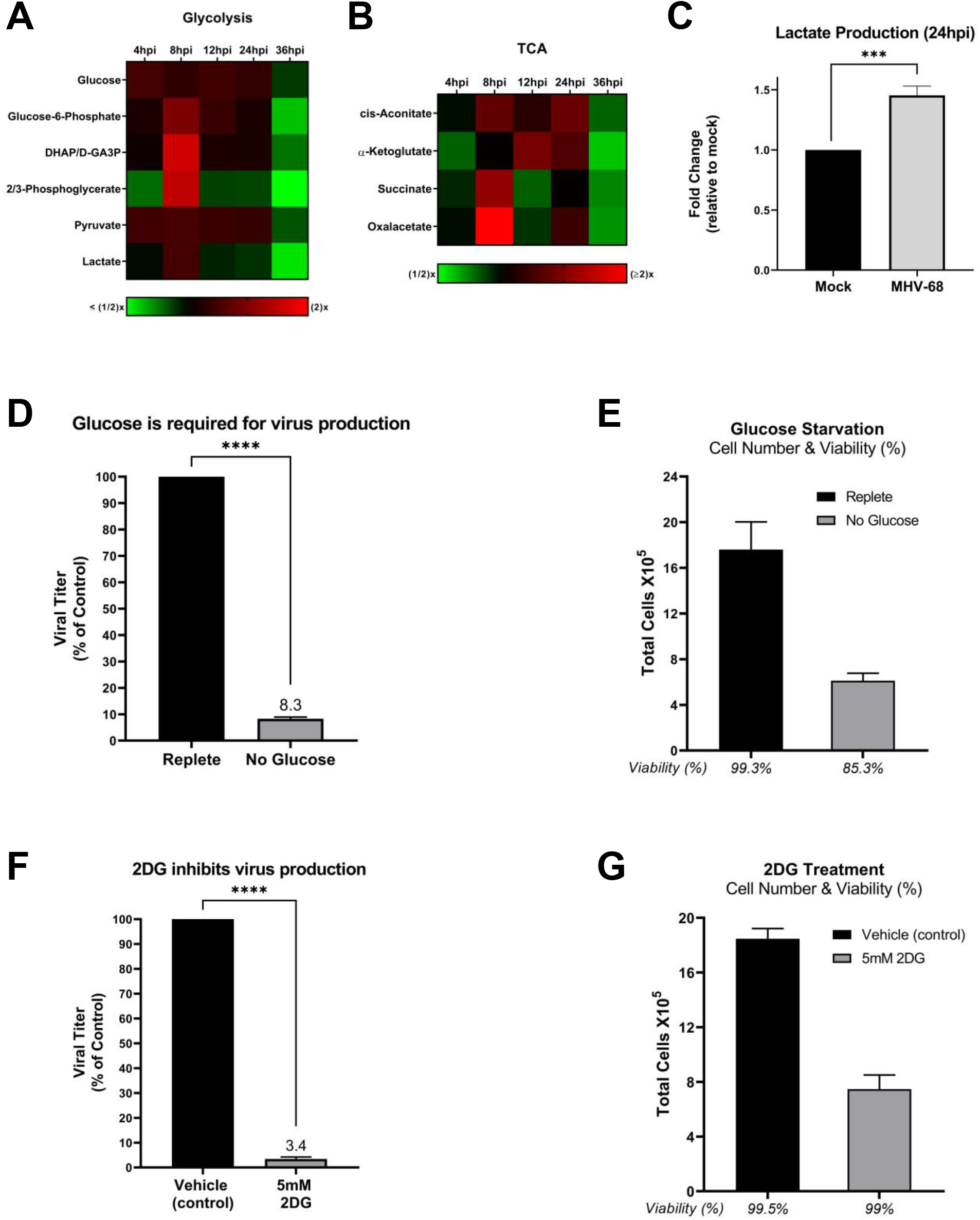
MHV-68 alters glucose metabolism and requires glucose for infectious virion production. A & B) Mock (control) and MHV-68 infected NIH3T3 cells were harvested at 4, 8, 12, 24, and 36 hpi in three independent experiments. Relative fold change time course for glycolytic and TCA cycle intermediates shown, where red depicts an increase and green shows a decrease in MHV-68 infected cells compared to mock infected cells. C) Lactate production is increased in MHV-68 infected NIH3T3 cells compared to mock infected (control) cells. Cellular supernatant was collected at 24 hpi and lactate production was quantified using the Promega Lactate-Glo assay kit in three independent experiments. Data was normalized to cell number. D & E) NIH3T3 cells were MHV-68 infected (MOI = 0.1) in three independent experiments. Cells were fed replete media (containing glucose and glutamine) for 24 hours and then replaced with fresh replete (control) or glucose-free media for an additional 24 hours. Viral titer was quantified from cell supernatants via plaque assays and graphed as % of control (replete media). Cell number and viability was calculated, using cellular pellets, via trypan blue exclusion assays with the Biorad TC20 automated cell counter. Cell number x 10^4 is graphed on the y-axis and viability (%) is listed under the x-axis for each condition. F & G) NIH3T3 cells were MHV-68 infected (MOI = 0.1) and treated with 5mM 2-deoxy-D-glucose (2DG) or vehicle (control) for 48 hpi in three independent experiments. Viral titer was quantified from cell supernatants via plaque assays and graphed as % of control (vehicle treatment). Cell number and viability was calculated, using cellular pellets, via trypan blue exclusion assays with the Biorad TC20 automated cell counter. Cell number x 10^4 is graphed on the y-axis and viability (%) is listed under the x-axis for each condition.

To determine the overall glycolytic output in MHV-68 infected cells vs mock infected cells, we measured the amount of lactate produced in cell supernatants at 24 hpi. Lactate is traditionally produced as a final product of glycolysis during anaerobic respiration, where pyruvate is converted to lactate and secreted outside the cell. However, recent data shows viral infections commonly increase lactate production in aerobic conditions in combination with an increase in glycolysis (7, 17, 39). Mock and MHV-68 infected cells (MOI = 3) were harvested at 24 hpi and lactate production was assessed in the cellular supernatant using an enzymatic assay. The data were normalized to the total cell number for each sample. As seen in figure 3C, there is a ∼50% increase in lactate production in MHV-68 infected cells compared to mock infected cells. Taken together, these data indicate MHV-68 infected cells increase glycolysis during lytic infection.

### MHV-68 lytic replication requires glucose for maximal virus production

To assess if glucose, the main carbon source for cells, is important for MHV-68 virion production, we quantified viral production at 48 hpi (one to two MHV-68 replication cycles) in the presence or absence of glucose. More specifically, mock and MHV-68 infected cells (MOI = 0.1) were first fed replete medium (containing glucose and glutamine) for 24 hours to minimize glucose starvation induced cell death. After 24 hours, the replete media was replaced with fresh replete medium (control) or glucose-free medium for an additional 24 hours. At 48 hpi, viral titers from MHV-68 infected cell supernatants were determined via plaque assays and plaque forming units (pfu) per mL were calculated. Cell viability and cell number of both mock and MHV-68 infected cells were determined via trypan blue exclusion assays. Our data show a ∼12-fold decrease in viral production in glucose-starved cells compared to cells treated with replete media (Fig. 3D). Glucose starvation during the final 24 hours of treatment resulted in a drop of cell viability to 85% compared to replete media fed cells which were 99% viable (Fig. 3E). Similar viability rates were seen in mock infected cells grown in replete or glucose-free media (data not shown). While there was a ∼3 fold decrease in cell number in glucose-starved cells compared to those in replete media (Fig. 3E), the drop in viral titer was more than 3-fold, indicating glucose is required for virus production.

To assess the need of glycolysis in viral production, we treated cells with the glycolytic inhibitor 2-deoxy-D-glucose (2DG). 2DG is a glucose analog which attenuates glycolysis by competing for the active site of hexokinase, which normally converts glucose to glucose-6-phosphate. To test if interfering with glycolysis results in decreased virus production, mock or MHV-68 infected cells (MOI = 0.1) were treated with vehicle (solvent control) or 5mM 2DG for 48 hours. Viral titers from MHV-68 infected cell supernatants were quantified via plaque assays and plaque forming units (pfu) per mL were calculated. Cell viability and cell number of both mock and MHV-68 infected cells were determined via trypan blue exclusion assays. Our data show a ∼30-fold decrease in viral production in 5mM 2DG treated cells compared to control treated cells (Fig. 3F). We also observed 48 hours of 2DG treatment did not significantly change infected cell viability compared to vehicle control treated cells (Fig. 3G). Similar viability rates were observed in mock infected cells treated with vehicle control or 2DG (data not shown). While there was a ∼3 fold decrease in cell number in 2DG treated infected cells compared to control, the drop in viral titer was ∼30 fold. Overall these data indicate glucose and increased glycolysis are important for viral production.

### MHV-68 lytic infection increases glutamine metabolism

Glutamine is an alternate carbon source to glucose. During glutaminolysis, glutamine is taken in by cells and converted to glutamic acid by the enzyme glutaminase (GLS). Glutamic acid is next converted into α-ketoglutarate by the enzyme glutamate dehydrogenase (GDH), which then enters the mitochondria. Overall, glutamine metabolism can fuel and replenish the TCA cycle, ultimately supporting increased ATP production and *de novo* lipid synthesis (lipogenesis). To determine if MHV-68 infection increases glutamine metabolism, we analyzed our aqueous metabolomics data for the abundance of metabolites in the glutaminolysis pathway (glutamine and glutamic acid). Fold change was calculated as a ratio of metabolite abundance in MHV-68 infected cells compared to mock infected cells. Our metabolomics analysis revealed MHV-68 infection increased both glutamine and glutamic acid metabolites at 4, 8 and 12 hpi (Fig. 4A). Next, we measured glutamine uptake in mock vs MHV-68 infected cells (MOI = 3). Mock and MHV-68 infected cells were harvested at 24 hpi and the amount of glutamine consumption was assessed in the cellular supernatant using an enzymatic assay. The data were normalized to total cell number. As seen in figure 4B, there is a ∼60% increase in glutamine consumption in MHV-68 infected cells compared to mock infected cells. We then determined the expression of glutaminolysis enzymes GLS and GDH in mock vs MHV-68 infected cells (MOI = 3) at 24 hpi. Western blot analysis showed no change in GLS expression (data not shown), but did show a ∼40% increase in GDH expression in MHV-68 infected cells compared to mock infected cells after densitometry analysis (Fig. 4C). Taken together, these data indicate MHV-68 infected cells increase glutamine metabolism during lytic infection.

**Figure 4:**
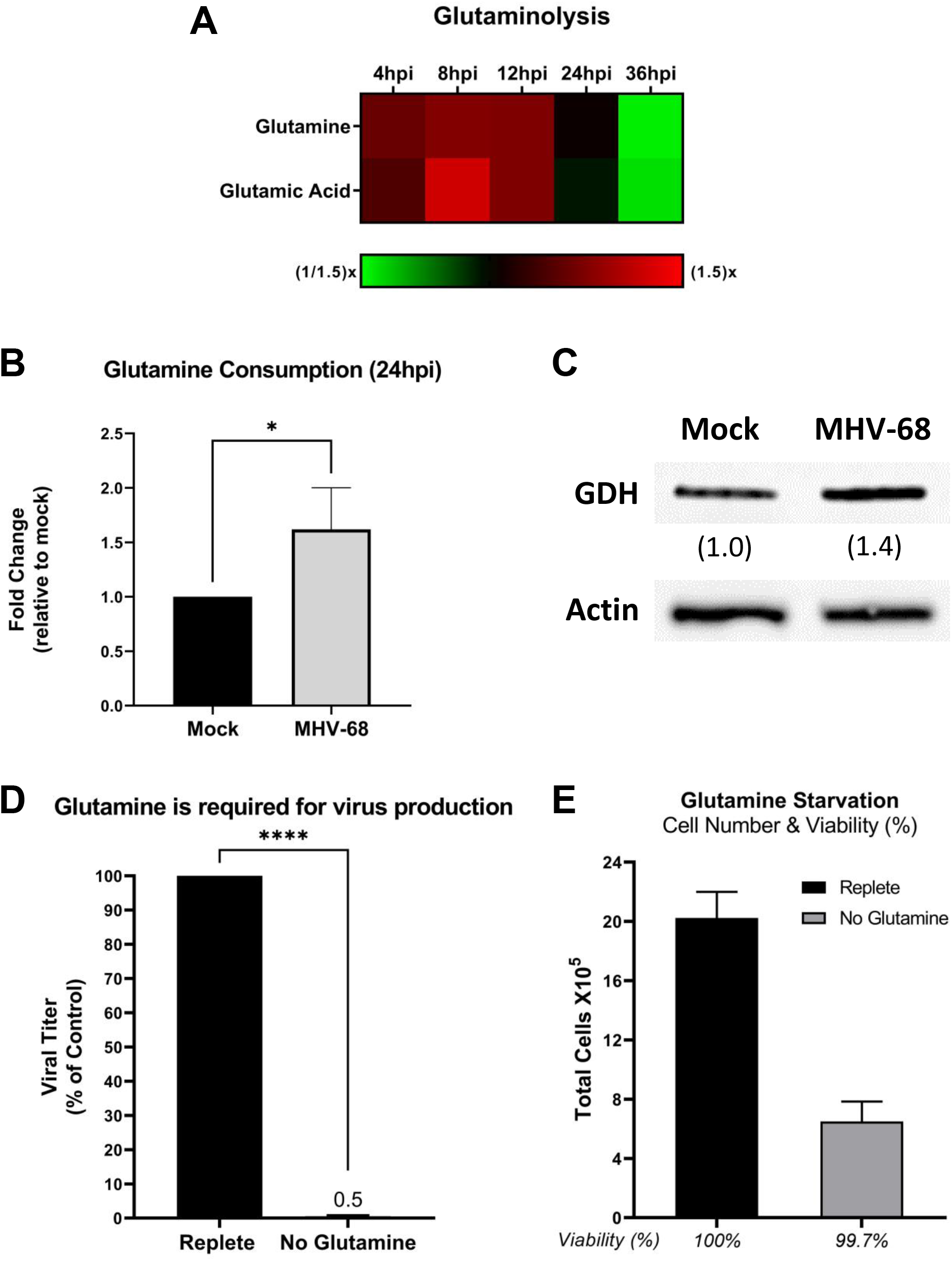
MHV-68 alters glutamine metabolism and requires glutamine for infectious virion production. A) Mock (control) and MHV-68 infected NIH3T3 cells were harvested at 4, 8, 12, 24 hpi, and 36hpi in three independent experiments. Relative fold change time course for glutaminolysis intermediates shown, where red depicts an increase and green shows a decrease in MHV-68 infected cells compared to mock infected cells. B) Glutamine consumption is increased in MHV-68 infected NIH3T3 cells compared to mock infected (control) cells. Cellular supernatant was collected at 24 hpi and glutamine consumption was quantified using the Promega Glutamine-Glo assay kit in three independent experiments. Data was normalized to cell number. C) Western blot analysis of glutamine dehydrogenase in mock vs MHV-68 infected (MOI = 3) NIH3T3 cells at 24 hpi in two independent experiments. Densitometry analysis shown as fold change compared to mock infected cells, using actin as a loading control. D & E) NIH3T3 cells were MHV-68 infected (MOI = 0.1) in three independent experiments. Cells were fed replete media (containing glucose and glutamine) or glutamine-free media for 48 hours. Viral titer was quantified from cell supernatants via plaque assays and graphed as % of control (replete media). Cell number and viability was calculated, using cellular pellets, via trypan blue exclusion assays with the Biorad TC20 automated cell counter. Cell number x 10^4 is graphed on the y-axis and viability (%) is listed under the x-axis for each condition.

### Glutamine is important for MHV-68 virion production

To assess the importance of glutamine for viral production, mock or MHV-68 infected cells (MOI = 0.1) were fed replete medium (containing glucose and glutamine) or glutamine-free medium for 48 hours. Viral titers from MHV-68 infected cell supernatants were determined via plaque assays and plaque forming units (pfu) per mL were calculated. Cell viability and cell number of both mock and MHV-68 infected cells were determined via trypan blue exclusion assays. Our data show a ∼190-fold decrease in viral production in glutamine-starved cells compared to cells fed replete media (Fig. 4D). Glutamine starvation did not significantly change infected cell viability compared to cells fed replete media (Fig. 4E). Similar viability rates were seen in mock infected cells fed replete or glutamine-free media (data not shown). While there was a ∼3 fold decrease in cell number in glutamine-starved infected cells compared to replete media fed cells (Fig. 4E), the drop in viral titer was ∼190-fold. Overall, glutamine starvation revealed glutamine is essential for MHV-68 production to an even higher degree than glucose starvation.

### Lipidomic profiling of lytic gammaherpesvirus infection reveals lipid remodelling throughout the viral life cycle

Lipids are a complex class of biological molecules which are important components of cell membranes and sources of cellular energy. Lipidomics is a bulk and quantitative analysis of lipids in cell samples which allows us to understand changes in lipid metabolism by creating a lipid profile of major lipid classes, subclasses, and individual species (40). To profile the host cell lipidome during lytic MHV-68 infection, we harvested mock or MHV-68 infected NIH3T3 cells (MOI = 3) for targeted lipidomics analysis at 4, 8, 12, 24 and 36 hpi in three independent experiments. Cells were harvested in -80°C cold methanol and sent for targeted lipidomics profiling at the University of Washington Northwest Metabolomics Research Center (NW-MRC) using the Sciex Lipidyzer mass spectrometry platform. Metabolite concentrations were normalized by whole cell protein concentration for each sample. Fold change was calculated as a ratio of lipid species abundance in MHV-68 infected cells compared to mock infected cells.

Our lipidomics analysis detected ∼300 unique lipid species at some point during the course of infection (Table S2). The category of lipid species detected included Cholesterol Ester (CE), Ceramides (CER), Diacylglycerol (DAG), Dihydroceramides (DCER), Free fatty Acids (FFA), Hexosylceramides (HCER), Lysophosphatidylcholine (LPC), Lysophosphatidylethanolamine (LPE), Phosphatidylcholine (PC), Phosphatidylethanolamine (PE), Sphingomyelin (SM), and Triacylglycerol (TAG). A detailed lipidomics analysis at 8 hpi, which detected 183 lipid species, revealed MHV-68 infected cells increased the abundance of 177 out of 183 (97% of detected lipids) and decreased the abundance of 54 out of 183 (3% of detected lipids) compared to mock infected cells (Figure 5A and Table S2). We next used volcano plot analysis to uncover the top statistically significant lipid species metabolites at 8 hpi (t-test, p<0.05, and at least 1.8 fold change). This analysis highlighted 22 significantly elevated lipid species (red) and 0 significantly decreased lipid species (Fig. 5B). Of these 22 elevated lipid species, 20 were TAG species and 2 were CE lipid species. CEs are dietary lipids composed of a cholesterol molecule, fatty acids, and a hydroxyl group. TAGs, which are composed of three fatty acid tails and a glycerol backbone, can be hydrolyzed to release free fatty acids, which are a major source of energy for cells.

**Figure 5:**
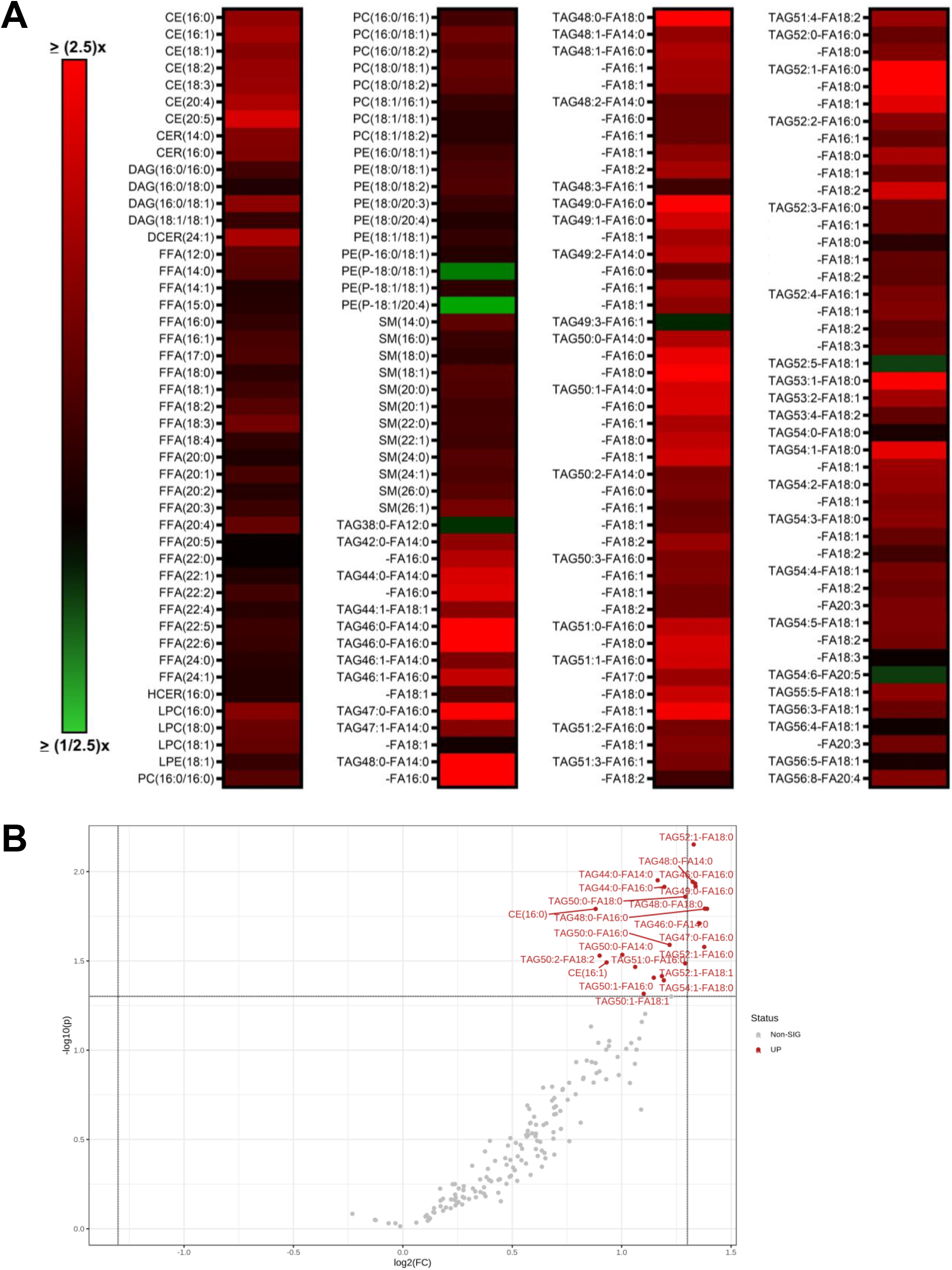
Quantitative lipidomics of mock vs MHV-68 infected cells at 8hpi. Mock (control) and MHV-68 infected NIH3T3 cells were harvested at 8 hpi in three independent experiments. CE = Cholesterol Ester, CER = Ceramides, DAG = Diacylglycerol, DCER = Dihydroceramides, FFA = Free fatty Acids, HCER = Hexosylceramides, LPC = Lysophosphatidylcholine, LPE = Lysophosphatidylethanolamine, PC = Phosphatidylcholine, PE = Phosphatidylethanolamine, SM = Sphingomyelin, and TAG = Triacylglycerol. A) Relative fold change depicted, where red shows an increase and green shows a decrease in lipid concentration in MHV-68 infected cells compared to mock infected cells. B) Volcano plot depicting log 2 fold change (FC) and -log10 p-value (p) in 8 hpi samples (paired). Statistical threshold set to p < 0.05 and at least 1.8 fold change in MHV-68 infected cells compared to mock infected cells using the student t-test. Red metabolites are significantly up.

Next, we analyzed our lipidomics data at 12 hpi, which detected 211 lipid species, and it revealed MHV-68 infected cells increased the abundance of 157 out of 211 (74% of detected lipids) and decreased the abundance of 54 out of 211 (26% detected lipids) compared to mock infected cells (Fig. 6A and Table S2). We next used the student t-test analysis to identify the top 4 most significant lipid species at 12 hpi (p<0.05, paired, and at least 1.5 fold change). Our analysis revealed a 1.9-fold increase in DAG (16:0 /18:0), a 1.6-fold increase in TAG55:5-FA18:1, a 1.6-fold increase in FFA (16:0), and a 1.5-fold increase in TAG56:3-FA18:1 in MHV-68 infected cells compared to mock infected cells (Fig. 6B and Table S2). To evaluate the changes in each lipid class over the course of infection, we graphed the pooled concentrations of each lipid class at 4, 8, 12 and 24 hpi (Fig. S1). TAG, FFA, and DAG lipid classes in mock vs MHV-68 infected cells over the course of infection revealed overall increases in TAGs after 8 hpi, increasing and peaking FFAs at 12 hpi, and increasing and peaking DAGs at 8-12 hpi before decreasing in concentration (Fig. 6C). Interestingly, all three of these lipid species are part of the same metabolism pathway. FFAs can be generated as 16-carbon fatty acids by lipogenesis and then elongated, or FFAs can be generated by the catabolic lipolysis of TAGs. DAGs are a glycerol molecule linked to two fatty acids and can be generated also by the creation or breakdown of TAGs. Overall, this data points to an early increase in TAG lipids (8 hpi) and then an increase in FAAs and DAG at 12 hpi, either by increased dietary intake of TAGs and subsequent lipolysis, lipogenesis, or both.

**Figure 6:**
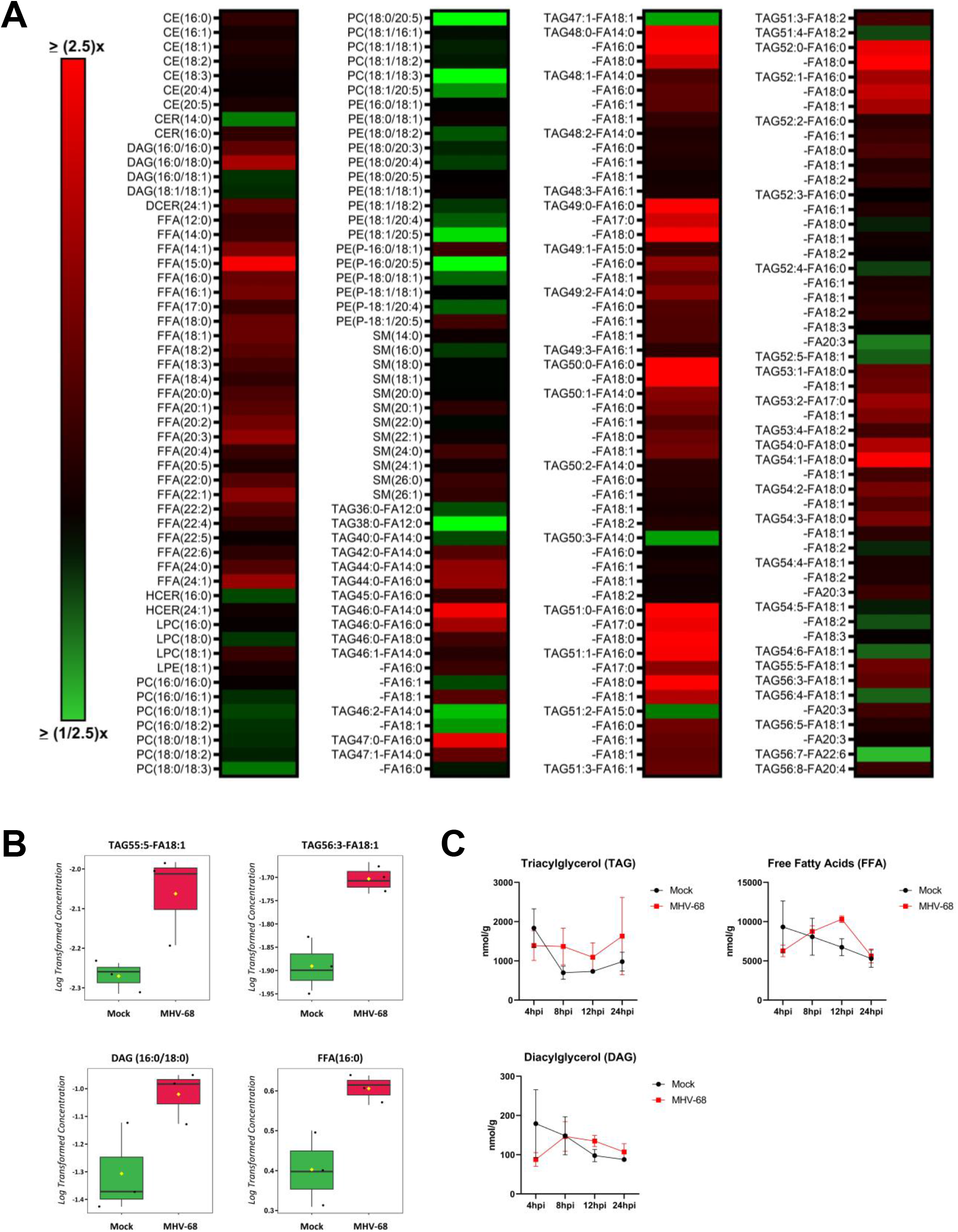
Quantitative lipidomics of mock vs MHV-68 infected cells at 12hpi. Mock (control) and MHV-68 infected NIH3T3 cells were harvested at 4, 8, 12, 24 or 36 hpi in three independent experiments. CE = Cholesterol Ester, CER = Ceramides, DAG = Diacylglycerol, DCER = Dihydroceramides, FFA = Free fatty Acids, HCER = Hexosylceramides, LPC = Lysophosphatidylcholine, LPE = Lysophosphatidylethanolamine, PC = Phosphatidylcholine, PE = Phosphatidylethanolamine, SM = Sphingomyelin, and TAG = Triacylglycerol. A) Relative fold change depicted, where red shows an increase and green shows a decrease in lipid concentration in MHV-68 infected cells compared to mock infected cells. B) Box plots of the top four statistically significant metabolites (t-test, p<0.05, and at least 1.5 fold change). Black dots represent concentrations for all samples (biological triplicates).The notch indicates the median and yellow diamond indicates the mean concentration. C) Total Lipid class concentration (TAG, FFA, and DAG) time course in mock (black circles) vs MHV-68 infected (red squares) NIH3T3 cells. Concentration (nmol/g) is shown on the y-axis and hours post infection on the x-axis.

### MHV-68 lytic infection increases fatty acid metabolism

*De novo* fatty acid synthesis (lipogenesis) results in the generation of new fatty acids and lipid materials. During lipogenesis, citrate is exported from the TCA cycle in the mitochondria and moves into the cytoplasm where it is converted to cytoplasmic acetyl-CoA by ATP citrate lyase (ACLY) and then malonyl-CoA by acetyl-CoA carboxylase (ACC). Both Acetyl-CoA and malonyl-CoA combine to form saturated 16-carbon fatty acid palmitic acid by fatty acid synthase (FASN), which is then used to make longer chain fatty acids and lipid materials. The induction of lipogenic enzymes are a key indicator of increased fatty acid synthesis in cells. Our western blot analysis of mock or MHV-68 infected cells (MOI = 3) at 24 hpi showed an increase in the lipogenic enzymes FASN and ACLY. Densitometry analysis of western blot data showed an ∼6.5-fold increase in FASN expression (Fig. 7A) and ∼2.25 fold increase in ACYL expression (Fig. 7B). There was no change in ACC expression (Fig. 7C). We analyzed our lipidomics data for the abundance of individual free fatty acids at 12 hpi (Fig. 7D and Fig. S2). Fold change was calculated as a ratio of metabolite abundance in MHV-68 infected cells compared to mock infected cells. Our analysis revealed that MHV-68 infection (MOI = 3) increased the abundance of all saturated FFAs detected by this lipidomics screen (Fig. 7D). MHV-68 infection also increased the abundance of all unsaturated fatty acids in our lipidomics analysis (Fig. S2). 16-carbon saturated FFAs showed the most significant change with a 1.6-fold increase in MHV-68 infected cells compared to mock infected cells at 12 hpi (p < 0.01). Taken together with our lipidomics data which shows overall increases in most lipid classes throughout the virus life cycle (Fig. S1), these data suggest both an increase in lipogenesis as well as dietary TAG uptake and lipolysis in cells with active MHV-68 lytic infection.

**Figure 7:**
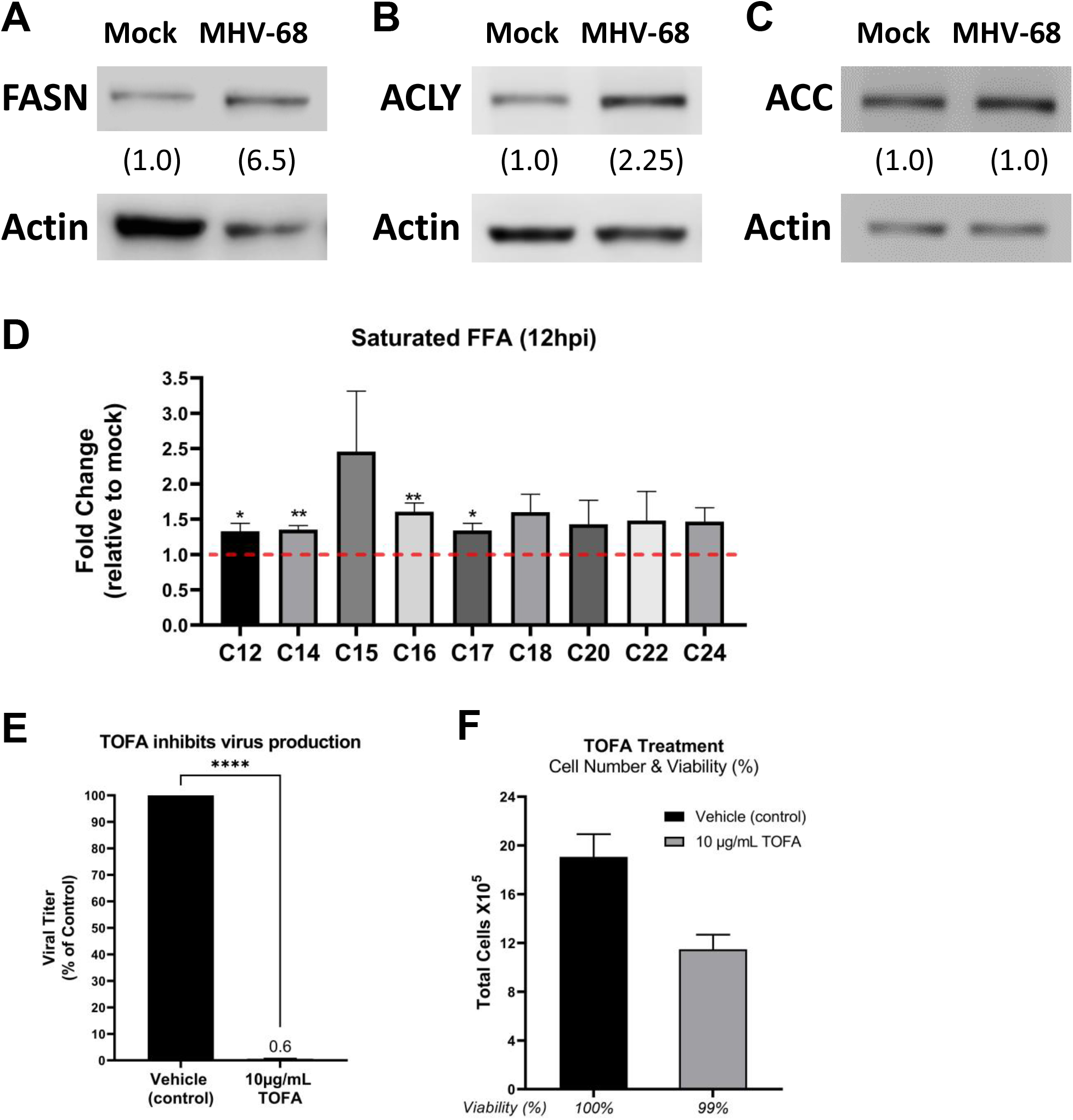
MHV-68 alters and requires lipogenesis for infectious virion production. (A, B, & C) Western blot analysis of Fatty Acid Synthase (FASN), ATP-Citrate Lyase (ACLY), and Acetyl-CoA Carboxylase (ACC), in mock vs MHV-68 infected (MOI = 3) NIH3T3 cells at 24 hpi in at least two independent experiments. Densitometry analysis shown as fold change compared to mock infected cells, using actin as a loading control. D) Quantitative lipidomics of saturated free fatty acids (FAA) in mock (control) and MHV-68 infected NIH3T3 cells at 12 hpi (three independent experiments). Unpaired t-test (p<0.05) data graphed on y-axis as relative fold change over mock. Red dashed line at 1.0 indicates no change compared to mock infected cells. E & F) NIH3T3 cells were MHV-68 infected (MOI = 0.1) and treated with 10 μg/mL TOFA or vehicle (control) for 48 hpi in three independent experiments. Viral titer was quantified from cell supernatants via plaque assays and graphed as % of control (vehicle treatment). Cell number and viability was calculated, using cellular pellets, via trypan blue exclusion assays with the Biorad TC20 automated cell counter. Cell number x 10^4 is graphed on the y-axis and viability (%) is listed under the x-axis for each condition.

### Lipogenesis is required for maximal virion production during lytic MHV-68 infection

The use of lipogenic inhibitors has been shown to block replication of several viruses, demonstrating these metabolic alterations are an important part of the viral life cycle (20, 41–44). 5-(Tetradecyloxy)-2-furoic acid (TOFA) is a pharmacological drug that inhibits the enzyme ACC in the lipogenesis metabolism pathway. To test if induction of lipogenesis is necessary for MHV-68 virion production, we mock or MHV-68 infected cells (MOI = 0.1) and treated them with vehicle control or 10 μg/mL TOFA. Viral titers from MHV-68 infected cell supernatants were determined via plaque assays and plaque forming units (pfu) per mL were calculated. Cell viability and cell number of both mock and MHV-68 infected cells were determined via trypan blue exclusion assays. Our data show a ∼160-fold log decrease in viral production in 10 μg/mL TOFA treated cells compared to control (Fig. 7E). TOFA treatment did not significantly change infected cell viability compared to control treated cells (Fig. 7F). Similar viability rates were seen in mock infected cells treated with solvent control or TOFA (data not shown). While there was a ∼1.7-fold decrease in cell number in TOFA treated cells compared to control, the drop in viral titer was ∼160-fold (Fig 7F). These data suggest that induction of lipogenesis driven by MHV-68 is necessary for virion production and fatty acid synthesis inhibitors are potential targets for gammaherpesvirus infections.

## DISCUSSION

While metabolomics analysis of host cells during viral infection has been investigated in multiple viruses (13, 15, 18, 30–32, 45–47), this study comprehensively analyzed changes in the host cell metabolome during gammaherpesvirus lytic replication. Our targeted aqueous metabolomics analysis revealed lytic MHV-68 infection induces many metabolic pathways, including glucose, glutamine, lipid, and nucleotide metabolism (Fig. 8). When we analyzed the top 12 statistically significant metabolites at 24 hpi, we discovered that 9 of the top 12 metabolites were nucleotide related species. The gammaherpesviruses KSHV, EBV, and MHV-68 all encode their own thymidine kinase, which aids in the synthesis of *de novo* DTMP and ultimately supports robust nucleotide production for viral replication (48–50). Not surprisingly, the top elevated metabolite in our metabolomics screen was DTMP. Other significant metabolites included the nucleotide cytosine, and related nucleosides cytidine and 2’-deoxy-cytidine, key building blocks in the creation of DNA and RNA. More recently, cytidine metabolism was revealed to be important during EBV virus induced B cell transformation, growth, and survival (51), highlighting its importance in gammaherpesvirus induced oncogenesis. The purine nucleoside 1-methyladenosine showed a significant increase during MHV-68 infection. Interestingly, during HIV-1 infection, it was revealed that HIV-1 virions contain abundant amounts of 1-methyladenosine, but this modification was shown to be linked solely to viral packaged tRNAs and not viral RNA and used to prime viral reverse transcriptase (52, 53). While gammaherpesviruses are DNA viruses and not retroviruses, it would be interesting to explore the role of 1-methyladenosine during gammaherpesvirus infection. Metabolomics analysis of two different herpesviruses, HCMV which has a ∼96 hour life cycle and HSV-1 which has a ∼24 hour life cycle, revealed divergent effects on cellular metabolism (13). While HSV-1 increased metabolic influx through the TCA cycle and pyrimidine nucleotide biosynthesis, HCMV increased glycolysis and the TCA cycle to support fatty acid synthesis. Due to the evidence that viruses trigger distinct metabolome changes in their host cells, it is important to metabolically profile viral families and individual viruses to understand which points in cellular metabolism would provide the best therapeutic targets.

**Figure 8:**
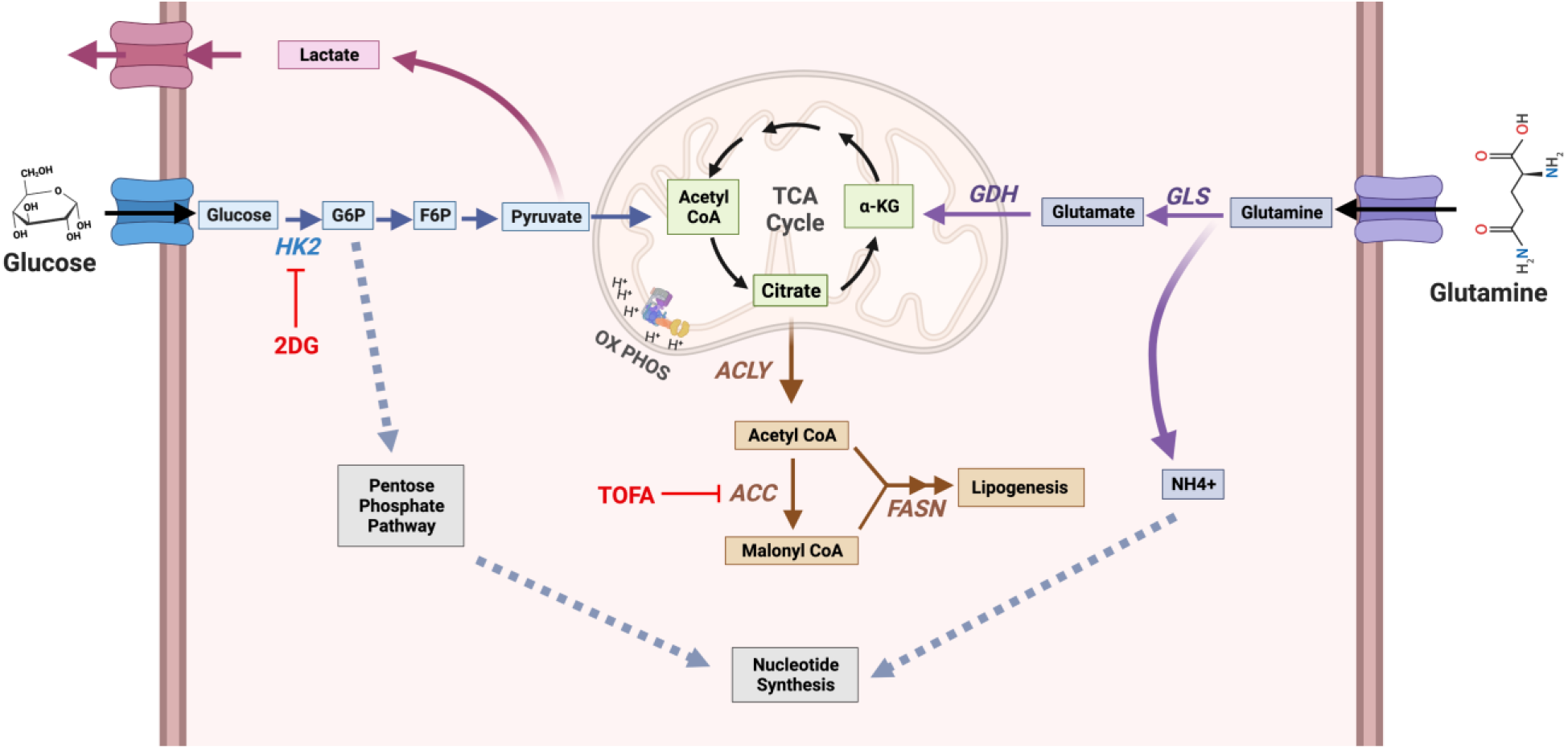
Model of host cell metabolism changes during lytic gammaherpesvirus infection. Our global metabolomics and lipidomics indicate that during MHV-68 lytic infection of host NIH3T3 cells, metabolic intermediates found in glycolysis, glutaminolysis, lipogenesis, and nucleotide metabolism, are significantly elevated over mock-infected controls. Briefly, glucose is taken up by the host cell via membrane bound glucose transporter proteins and metabolized by Hexokinase (HK) to form glucose-6-phosphate (G-6-P). Depending on cellular or viral need, G-6-P can have several fates, including 1) be further metabolized through glycolysis to form pyruvate to fuel the TCA cycle in the mitochondria, 2) be converted from pyruvate to lactate and secreted outside of the cell, or 3) fuel the pentose phosphate pathway for nucleotide synthesis. During glutaminolysis, glutamine is taken up by the host cell via a membrane bound glutamine transporter. Glutamine is converted into glutamate and NH_3_ by glutaminase (GLS), and then α-ketoglutarate by glutamate dehydrogenase (GDH). Next, α-ketoglutarate enters and replenishes the TCA cycle.The NH_3_ generated from glutamine breakdown can be used for nucleotide synthesis. Overall, downstream intermediates from both glucose and glutamine, which feed into the TCA cycle, can fuel the lipogenesis pathway. Briefly, citrate is converted to cytoplasmic acetyl-CoA by ATP citrate lyase (ACLY) and then malonyl-CoA by Acetyl-CoA carboxylase (ACC). Both Acetyl-CoA and malonyl-CoA combine to form palmitic acid by Fatty Acid Synthase (FASN), which is then used as a backbone to make longer chain fatty acids, triglycerides, phospholipids, and other lipid materials. During MHV-68 lytic infection, key enzymes in these pathways, including GDH, ACLY, and FASN are elevated in expression.

Glucose, the main central carbon source of cells, feeds into glycolysis and subsequently fuels cellular ATP production. Glucose is also an important carbon source to fuel other metabolic pathways such as the pentose phosphate pathways for nucleotide production and lipogenesis through the TCA cycle for *de novo* fat production (Fig. 8) (54). The catabolism of glutamine, an alternate carbon source to glucose, replenishes the TCA cycle. This is particularly important when the TCA cycle is depleted due to glucose being metabolized to lactate instead of pyruvate, and citrate being shuttled out of the TCA cycle to support fatty acid synthesis (Fig. 8). Therefore, glutamine breakdown and fueling of the TCA cycle ultimately supports increased ATP production and lipid synthesis (Fig 8). Glutamine breakdown also provides an important nitrogen source for nucleotide synthesis (55, 56). Interestingly, increased glucose and glutamine metabolism have been shown to be key hallmarks of cancer and are often required for induction of lipogenesis (57, 58). We and others have previously investigated whether glucose or glutamine metabolism are induced by infection or required for viral production. For example, we showed latent KSHV infection requires both glucose and glutamine for the maintenance of latently infected endothelial cells (17, 19). Furthermore, we showed that induction of lytic replication during KSHV infection was reduced when host cells were starved of glucose or glutamine, with glucose being more important than glutamine for virion production (20). However, studies in HCMV and VAVC show glutamine but not glucose is required for infectious virus production (15, 59). These studies reveal that the need for glucose or glutamine during infection is virus dependent, and understanding which viruses prefer glucose vs glutamine can help provide clues for more selective viral therapeutics.

Our metabolomics data show an increase in glycolytic and TCA intermediates at 8 hpi. We also show an increase in lactate production, an output of glycolysis, at 24 hpi. We and others have shown 2DG treatment, which attenuates glycolysis, inhibits MHV-68 production (29, 60), demonstrating glycolysis is required for viral production. Additionally, our metabolomics data show an increase in glutaminolysis intermediates at 4 hpi, 8 hpi, and 12 hpi. Furthermore, we observed an increase in glutamine consumption and glutamate dehydrogenase expression at 24 hpi, highlighting an overall increase in glutamine metabolism during MHV-68 lytic replication. Interestingly, we demonstrated that glucose starvation of host cells led to a ∼12 fold decrease in infectious virus production while glutamine starvation of host cells led to a ∼190 fold decrease in virus production. While our data demonstrate that MHV-68 infected cells increase and require both glucose and glutamine metabolism, our data strongly support that glutamine is more important than glucose for MHV-68 virion production. It is possible that glutamine may be necessary to feed multiple host cell metabolic pathways, including the TCA cycle for energy production and lipogenesis, and nucleotide metabolism for the synthesis of *de novo* DNA and RNA (Figure 8). Further labelled glutamine flux experiments are needed to track glutamine utilization to ultimately determine its role during MHV-68 lytic infection and its necessity during viral production.

Essential alterations in the cellular lipid profile have been observed for many tumor models (61–64). Induced lipid metabolism fills energy stores for the cell as well as serve as biosynthetic material for cellular membranes and other lipid intermediates. Lipid metabolism has been proven to be critical for tumorigenesis and tumor cell survival (63). Like human gammaherpesvirus, one of the most important implications of studying MHV-68 infection is understanding viral-induced oncogenesis and tumor formation. Our lipidomics data reveal a clear gammaherpesvirus lipid profile is established during lytic gammaherpesvirus infection. We show pharmacological inhibition of lipogenesis blocks virus production, indicating that gammaherpesvirus virion production is dependent on this modulation in lipid metabolism.

Previous work has shown that other herpesviruses also depend on alterations of lipid metabolism (13, 18, 20, 28, 65–68). Inhibition of lipogenesis, specifically synthesis of phosphatidylethanolamine (PE), has been shown to inhibit viral production and budding in HSV-1 (69). We see an increase in PE detection in our data at eight hours post MHV-68 infection. This suggests specific lipid requirements may be critical for herpesvirus dissemination. Additionally, HCMV induces adipocyte-like conditions by significantly inducing lipogenesis (65). Specifically, sterol regulatory element-binding transcription factor 1a (SREBP1a) and SREB1c are involved in mediating this adipocyte-like phenotype and the inhibition of SREBP1 leads to impaired HCMV growth. It is possible that SREBP1 contributes to the mechanism by which gammaherpesviruses induce lipogenesis, as we see a significant increase in numerous lipid species in our data. Finally, several RNA viruses also hijack host lipid metabolism, including Dengue virus, Hepatitis C virus, Influenza virus, and SARS-CoV-2 (44, 70, 71). Positive-single stranded RNA viruses enhance lipid droplet formation during early infection and hijack membranes during replication (72). Inhibiting lipid biosynthesis and formation of lipid droplets negatively impacts viral replication of these viruses. Exploring the role of lipid droplets and lipid droplet composition during lytic gammaherpesvirus infection could further inform whether these structures and their components are essential for viral replication and virion production.

Overall, previous work has shown that central carbon metabolism is essential for the replication of KSHV, a related gammaherpesvirus (20). We showed that inhibition of glycolysis, glutamine metabolism, or fatty acid synthesis blocked KSHV viral replication at distinct steps of viral gene transcription, translation, or virion assembly. While it is possible that additional metabolic alterations are required for maximal KSHV replication and virion production, a global analysis of metabolic changes was not included. Using MHV-68, a model gammaherpesvirus, our global metabolomics and lipidomics data demonstrate that gammaherpesvirus replication is highly dependent on induced central carbon metabolism. We show that glucose, glutamine, and lipid metabolism are essential for maximal infectious virus production, and inhibition of these pathways via nutrient depletion or drug inhibition results in a significant reduction of virus production. Identifying that these key metabolic pathways are critical for the production of infectious virions is the first step in selecting molecular targets for pharmacological inhibition. Our findings provide a framework for future *in vivo* experiments in which high priority metabolic inhibitors can be tested to block virus infection and cancer progression in mice infected with MHV-68. Ultimately, the translation of this mouse model system can be used to treat gammaherpesvirus induced cancers and infections in humans.

## MATERIALS AND METHODS

### Cell lines and reagents

NIH3T3 cells (ATCC #CRL-1658) or Vero cells (ATCC #CCL-81) were maintained at 37°C in 5% CO_2_ in as monolayer cultures in DMEM media with L-Glutamine and sodium pyruvate (Genesee # 25-500). NIH3T3 cell media was supplemented with 10% newborn calf serum (NBCS) and 1% penicillin streptomycin (P/S). Vero cell media was supplemented with 10% fetal bovine serum (FBS) and 1% P/S. 2M stocks of 2-deoxy-D-glucose (2DG) (Sigma # D6134) were diluted in dH_2_O and 5 mg/mL stocks of 5-(Tetradecyloxy)-2-furoic acid (TOFA) (SCBT # 200653) were diluted in DMSO.

### Viruses, infection, and plaque assays

To propagate virus stocks, NIH3T3 or Vero cells were infected with MHV-68 (ATCC #VR-1465) at a MOI of 0.01 – 0.05. When ∼80% of the cells detached from the plate, the cells and media were freeze thawed and pelleted at 3,000 rpm for 10 min at 4°C. The cleared viral supernatant was concentrated by centrifugation at 20,500 x g at 4°C for 90 min. Viral pellets were resuspended in serum free DMEM. Viral infections were performed at a multiplicity of infection (MOI) of 0.1 or 3 in DMEM for 2 hrs, after which the media was replaced with fresh DMEM. Viral titers were determined using traditional plaque assays. Briefly, four hours prior to infection, 12 well plates seeded with Vero cells (250,000 cells/well). Virus samples were serial diluted (10^0 - 10^6) in serum free (SF) DMEM and Vero cells were infected for 1.5 hours. Viral dilutions were then aspirated, and cells were overlayed with DMEM + glucose + glutamine + pyruvate (Thermofisher #12800017) + 1% methylcellulose (MC) (Sigma # M0387) + sodium bicarbonate (Sigma #S5761) + 2.5ug/mL Amphotericin B (Thermofisher # 15290026), 5% FBS + 1% P/S for 6 days. Cells were fixed in 10% formalin, media overlay was removed, cells were stained with 1% crystal violet (Sigma #C0775) in 20% methanol and rinsed with water before drying.

### Western blot analysis

Mock or MHV-68 infected NIH3T3 cells (MOI = 3) and media were harvested at 24 hours post infection (hpi) using a cell scraper and pelleted at 200 x g for 5 min at 4°C. Cell pellets were washed in cold PBS and centrifuged at 13000 rpm at 4°C. Cell pellets were resuspended in RIPA lysis buffer (Sigma #R0278) + protease inhibitor cocktail (Sigma # S8820) + phosphatase inhibitor cocktail (Sigma #524629) on ice for 30 min. Protein concentrations were determined by a Pierce BCA protein assay kit (Thermofisher # 23225). 10-20 ug of protein was separated by a sodium dodecyl sulfate polyacrylamide (SDS-PAGE) gel and transferred to a polyvinyldifluoride or nitrocellulose membrane, and blotted at the appropriate primary antibody [dilutions were 1:1,000 mouse anti-actin (SCBT #47778) or 1:2,500 rabbit anti-actin (∼43kDa) (Cell Signalling # 4970), 1:1,000 anti-glutamine dehydrogenase ½ (∼50-55 kDa) (SCBT # 515542), 1:500 anti-fatty acid synthase (∼270kDa) (SCBT # 20140), 1:1,000 anti-ATP-citrate lyase (∼125kDa) (Cell Signalling #4332), and subsequently with horseradish peroxidase (HRP)-conjugated 1:5,000 goat-anti-mouse (Licor #926-80010) or 1:5,000 goat-anti-rabbit (Licor # 926-80011), or infrared conjugated 1:10,000 goat-anti-mouse-680 (Licor # 926-68070) or 1:5,000 goat-anti-rabbit-800 (Licor # 926-32211). HRP-conjugated proteins were incubated in WesternSure Chemiluminescent substrate (Licor #926-95000). Immunoreactive proteins were detected using the Licor Odyssey Fc or C-DiGit imagers. Densitometry analysis of protein expression was determined using the Licor Image Studio or Licor Empiria Studio software, with actin set as the loading control.

### Glucose starvation, Glutamine starvation, or Drug assays

NIH3T3 cells were seeded at a density of 760,000 cells in 6 cm dishes approximately 4 hours prior to infection. MHV-68 infected cells were infected at a MOI of 0.1 for two hours. For both glucose and glutamine starvation assays, a base medium of DMEM (no glucose, no glutamine, no phenol red) (Thermofisher # 11960044) was used. Media was supplemented with 10% newborn calf serum, 1% penicillin and streptomycin, 2mM L-Glutamine (Genesee #25-509), and/or 4.5g/L glucose (Thermofisher # A2494001). After mock or MHV-68 infection, cells were washed 3x with PBS and then fed replete (glucose + glutamine), glucose-free (+ glutamine), or glutamine-free (+ glucose) media. During glucose starvation experiments, cells were fed replete media for 24 hours, washed 3X with PBS, and then the media was replaced with replete or glucose-free media for the final 24 hours. For 2DG or TOFA experiments, mock or MHV-68 infected cells were replaced with complete DMEM (containing 2mM glutamine, 1% penicillin-streptomycin and 10% newborn calf serum) and their respective drug treatment. 2DG was added to the cell medium at a final concentration of 5mM in DMSO. TOFA was added to cell culture medium at a final concentration of 10 µg/mL in water. Cell supernatants were cleared by centrifugation and used to determine viral titers via plaque assays and cell pellets were harvested to determine cell number and viability via trypan blue exclusion assays.

### Cell number and viability

Cell supernatants and trypsinized cells from 6 cm dishes were centrifuged for 5 min. Cell pellets were resuspended in 500uL of complete DMEM. Cell number and viability was determined via trypan blue exclusion assays using the Biorad TC20 automated cell counter (Biorad # 1450102). Briefly, 1:1 ratios of cells to trypan blue were mixed and loaded on dual chambered slides (Biorad # 1450003). Total cell numbers, live cell numbers, and percent viability were calculated by the Biorad TC20.

### Glutamine consumption and lactate production

NIH3T3 cells were seeded at a density of 700,000 cells in 6 cm dishes approximately 4 hours prior to infection. MHV-68 infected cells were infected at a MOI of 3 for two hours. Mock or MHV-68 infected cells were washed with PBS and replaced with DMEM (no glucose, no glutamine, no phenol red) (Thermofisher # 11960044) supplemented with 10% dialyzed fetal bovine serum (Thermofisher # A3382001), 1% penicillin and streptomycin, 2mM L-Glutamine (Thermofisher #25030149), and 5mM glucose (Thermofisher # A2494001). At 24 hpi, cell supernatants were cleared by centrifugation and cell pellets were harvested to determine cell number and viability via trypan blue exclusion assays using the Biorad TC20. Cell supernatants were assayed for glutamine consumption (Promega #J8021) and Lactate production (Promega #J5921) according to the kit specifications. Glutamine consumption and Lactate production was normalized to cell number.

### Aqueous Metabolomics and Lipidomics Experimental Set Up

NIH3T3 cells were seeded in 10 cm dishes (∼1.5 million cells) or 6 cm dishes (∼ 720,000 cells) approximately 4 hours prior to infection. MHV-68 infected cells were infected at a MOI of 3 for two hours. After infection, mock or MHV-68 infected cells were fed complete DMEM (containing 2mM glutamine, 1% penicillin-streptomycin, and 10% newborn calf serum). In biological triplicates at 4, 8, 12, 24 and 36 hpi, cell metabolism was quenched by harvesting and rinsing with -80°C chilled methanol on ice with cell scrapers. Cells were centrifuged for 5 min at 125 x g (4°C) and the supernatant was aspirated. Cell pellets were flash frozen in liquid nitrogen and transported to the University of Washington Northwest Metabolomics Research Center (NW-MRC) for analysis. The NW-MRC provided metabolic profiling of samples using targeted aqueous metabolomics and quantitative lipidomics platforms.

### Aqueous Metabolomics

<u>Sample Prep</u>: Aqueous metabolites for targeted liquid chromatography - mass spectrometry (LC-MS) profiling cell samples were extracted using a protein precipitation method similar to the one described elsewhere (73, 74). Samples were first homogenized in 200 µL purified deionized water at 4 °C, and then 800 µL of cold methanol containing 124 µM 6C13-glucose and 25.9 µM 2C13-glutamate was added (reference internal standards were added to the samples in order to monitor sample prep). Afterwards, samples were vortexed, stored for 30 minutes at -20 °C, sonicated in an ice bath for 10 minutes, centrifuged for 15 min at 14,000 rpm and 4 °C, and then 600 µL of supernatant was collected from each sample. The protein pellets were saved for BCA assay (total protein count was needed for data normalization). Lastly, recovered supernatants were dried on a SpeedVac and reconstituted in 1.0 mL of LC-matching solvent containing 17.8 µM 2C13-tyrosine and 39.2 3C13-lactate (reference internal standards were added to the reconstituting solvent in order to monitor LC-MS performance). Samples were transferred into LC vials and placed into a temperature controlled autosampler for LC-MS analysis. <u>LC-MS Assay</u>: Targeted LC-MS metabolite analysis was performed on a duplex-LC-MS system composed of two Shimadzu UPLC pumps, CTC Analytics PAL HTC-xt temperature-controlled auto-sampler and AB Sciex 6500+ Triple Quadrupole MS equipped with ESI ionization source (74). UPLC pumps were connected to the auto-sampler in parallel and were able to perform two chromatography separations independently from each other. Each sample was injected twice on two identical analytical columns (Waters XBridge BEH Amide XP) performing separations in hydrophilic interaction liquid chromatography (HILIC) mode. While one column was performing separation and MS data acquisition in ESI+ ionization mode, the other column was getting equilibrated for sample injection, chromatography separation and MS data acquisition in ESI-mode. Each chromatography separation was 18 minutes (total analysis time per sample was 36 minutes). MS data acquisition was performed in multiple-reaction-monitoring (MRM) mode. LC-MS system was controlled using AB Sciex Analyst 1.6.3 software. Measured MS peaks were integrated using AB Sciex MultiQuant 3.0.3 software. The LC-MS assay was targeting 361 metabolites (plus 4 spiked reference internal standards), and 172 metabolites (plus 4 spiked internal standards) were measured across the study set. In the addition to the study samples, two sets of quality control (QC) samples were used to monitor the assay performance as well as data reproducibility. One QC [QC(I)] was a pooled human serum sample used to monitor system performance and the other QC [QC(S)] was pooled study samples and this QC was used to monitor data reproducibility. Each QC sample was injected per every 10 study samples. The data were well reproducible with a median CV of 5.9 %.

### Quantitative Lipidomics

<u>Sample Prep</u>: 250 μL of water and 1 mL of cold methanol were added to frozen cell samples in Eppendorf tubes, vortexed for 10 seconds and the mixtures were transferred to glass culture tubes. 250 μL of water was added to previously emptied Eppendorf tubes, tubes were vortexed for 10 seconds and the content was transferred into the glass culture tubes to recover any cells that were left on the walls of the Eppendorf tubes. 450 μL of dichloromethane was added to the mixture in the glass tubes, vortexed for 10 seconds and 25 μL of a mixture containing 54 isotope labelled internal standard lipids was added to the tubes. Samples were sonicated in an ice batch for 10 min and then left to incubate at room temperature for 30 min. Next, another 500 μL of water and 450 μL of dichloromethane were added followed by vortexing for 10 s and centrifugation at 2500×g at 15 °C for 10 min. The bottom organic layer was transferred to a new tube and 900 μL of dichloromethane were added to the original tube for a second extraction. The combined extracts were concentrated under nitrogen and reconstituted in 250 μL of the running solution (10 mM ammonium acetate in 50:50 methanol:dichloromethane). At this point the samples were ready for mass spectrometry analysis. The protein precipitate that was left-over from sample prep was used for BCA assay (total protein content was needed for data normalization). <u>Mass Spectrometry Data Acquisition</u>: Lipid and fatty acid species in the extracted cell samples were measured as described elsewhere (75). The Sciex Lipidyzer mass spectrometry platform consisted of Shimadzu Nexera X2 LC-30AD pumps, a Shimadzu Nexera X2 SIL-30AC autosampler, and a Sciex QTRAP 5500 mass spectrometer equipped with SelexION for differential mobility spectrometry (DMS). 1-propanol was used as the chemical modifier for the DMS. Samples were introduced to the mass spectrometer by flow injection analysis at 8 uL/min. Each sample was injected twice, once with the DMS on (PC, PE, LPC, LPE and SM), and once with the DMS off (CE, CER, DAG, DCER, FFA, HCER, LCER and TAG). The lipid molecular species were measured using multiple reaction monitoring (MRM) and positive/negative polarity switching. SM, DAG, CE, CER, DCER, HCER, DCER and TAG were ionized in positive and LPE, LPC, PC, PE and FFA in negative ionization mode, respectively. A total of 1070 lipid species and fatty acids were targeted in the analysis. Data were acquired and processed using Analyst 1.6.3 and Lipidomics Workflow Manager 1.0.5.0. 205-471 lipids were measured across the study set of 30 samples. All the samples were prepared and analyzed in a single batch. For quality control, an instrument QC of pooled human serum was run at the beginning and at the end of the batch, respectively. The median CV was 4.5%.

### Metabolomics Data Analysis

For each metabolite (lipid or aqueous) concentration was divided by total protein to scale for the variable numbers of cells in each sample as follows, S=M/P, where S is the scaled value, M is the absolute lipid or aqueous metabolite concentration, and P is the protein concentration in the sample. For each time point on each run, the virally infected sample’s scaled value was divided by that of the corresponding control (i.e., mock infections) as follows, R=S_trv_/S_trm_, where R is the resulting ratio of virus scaled value: mock infection scale value for a given lipid or aqueous metabolite, S_trv_ is the scaled value for a given time point and run in the virally-infected replicate and S_trm_ is the equivalent value for the mock infection replicate. Mean R was found for each time point by averaging across all 3 runs (biological triplicate). Metabolomics heat maps were generated using Graphpad Prism 8 software as fold change, where red shows an increase and green shows a decrease in MHV-68 infected cells compared to mock infected cells. Volcano plots and box plots were generated using the freeware software MetaboAnalyst 5.0 (https://www.metaboanalyst.ca/home.xhtml). Normalized metabolite concentrations (mock vs MHV-68 infected cells) were uploaded on the Metaboanalyst website under the Statistical Analysis (one-factor) module. Data was analyzed in Metaboanalyst using the Student’s (two-tailed) t test. Graphs were generated as MHV-68/Mock, with p values set to 0.05 or lower and fold change cut offs of 1.5 fold or higher. Data was log transformed and volcano plot or box plots were downloaded for this publication. Volcano plots were graphed as log2 fold change (FC) on the x-axis and -log 2 (p) on the y-axis. For the box plot graphs, the black dots represent concentrations for all samples in biological triplicates. The notch indicates the median with 95% confidence, as defined as +/- 1.58*IQR/sqrt(n). The yellow diamond indicates the mean concentration.

### Statistical Analysis and Graphs

All bar and line graphs were generated using Graphpad Prism 8 software. Statistical analysis of bar graphs was performed using the Prism software. Standard errors of the mean (SEM) are shown and p-values were analyzed using the Student’s (two-tailed) t-test. A p-value of <0.05 is considered statistically significant and indicated by a asterisk (*). A p-value of <0.01 is indicated by two asterisk (**), a p-value of <0.001 is indicated by three asterisks (***), and a p-value of <0.0001 is indicated by four asterisks (****).

## Supporting information

Table S1 - Aqueous metabolomics normalized fold change time course in mock vs MHV-68 infected cells.

Table S2 - Quantitative lipidomics normalized fold change time course in mock vs MHV-68 infected cells.

## ACKNOWLEDGEMENTS

The Delgado lab metabolomics and lipidomics project was sponsored by a grant from the Supporting Structures: Innovative Partnerships to Enhance Bench Science at CCCU Member Institutions program, run by Scholarship and Christianity in Oxford, the UK subsidiary of the Council for Christian Colleges and Universities, with funding by the John Templeton Foundation and the MJ Murdock Charitable Trust. Other Delgado lab funding sources include the Murdock College Research Program for Natural Sciences grant, Seattle Pacific University Faculty Research and Scholarship Grants, and the Northwest University Ray and Shirley Clark Faculty Research Grant. Osvaldo Kevin Moreno was supported through a Genentech Foundation Scholarship. We would like to thank Dr. Michael Lagunoff for support. We also acknowledge the collaborative services of The Northwest Metabolomics Research Center at the University of Washington, Seattle and NIH S10 grant # 1S10OD021562-01 that funded a purchase of the Sciex mass spectrometry platform that was used to acquire metabolomics (quantitative lipidomics) data. We would finally like to thank Delgado lab undergraduate researchers Glenwood Ray Clark and Aubrey Massman for helping set up general lab protocols, and Tyler Ptacek and Anna Miller for setting up preliminary experiments not included in this manuscript.

---

**Figure S1:**
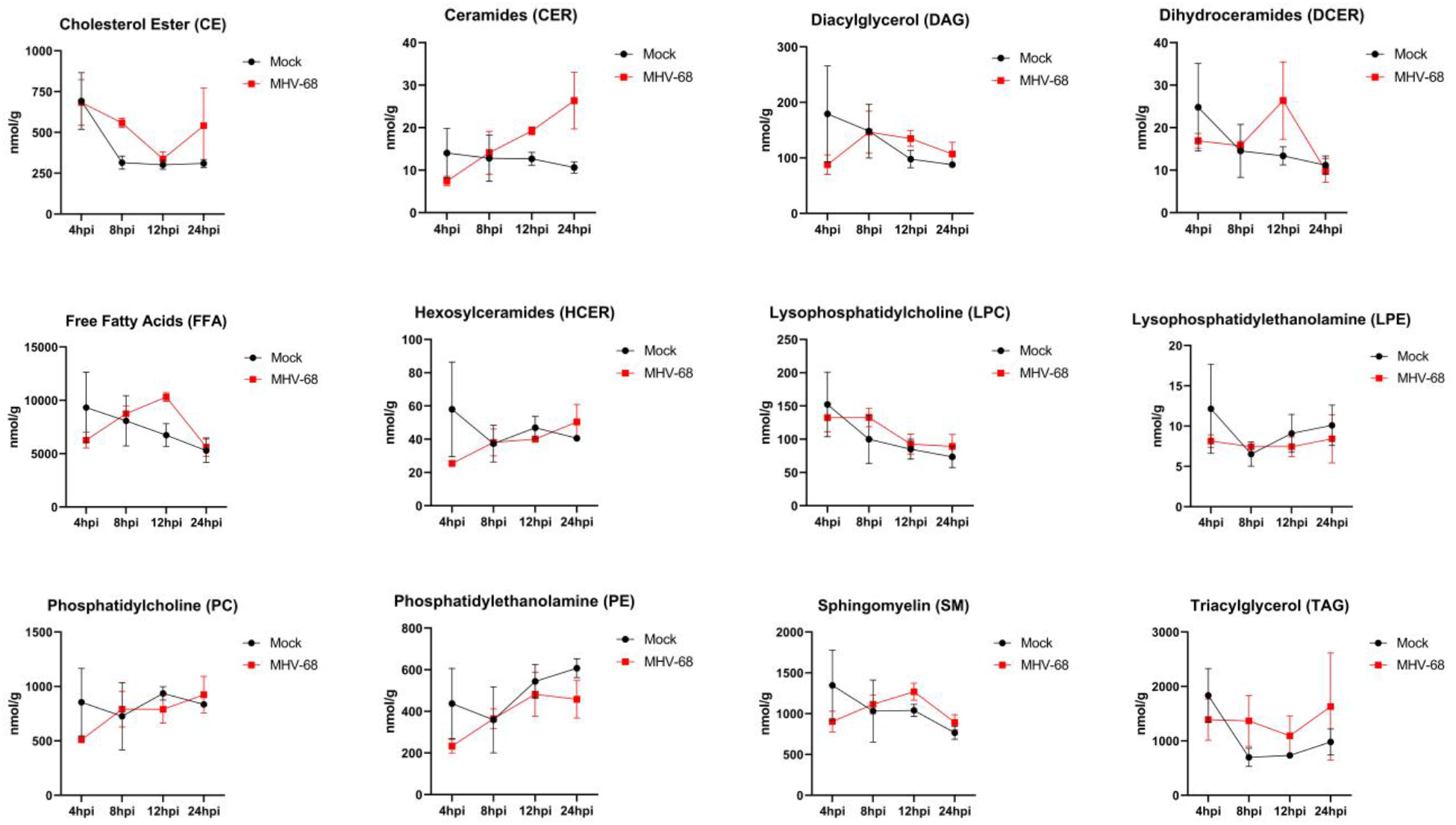
Total lipid class concentration time course of mock vs MHV-68 infected cells. Mock (control) and MHV-68 infected NIH3T3 cells were harvested at 4, 8, 12, 24 or 36 hpi in three independent experiments. CE = Cholesterol Ester, CER = Ceramides, DAG = Diacylglycerol, DCER = Dihydroceramides, FFA = Free fatty Acids, HCER = Hexosylceramides, LPC = Lysophosphatidylcholine, LPE = Lysophosphatidylethanolamine, PC = Phosphatidylcholine, PE = Phosphatidylethanolamine, SM = Sphingomyelin, and TAG = Triacylglycerol. Data shows total lipid class concentration time course in mock (black circles) vs MHV-68 infected (red squares) NIH3T3 cells. Concentration (nmol/g) is shown on the y-axis and hours post infection on the x-axis.

**Figure S2:**
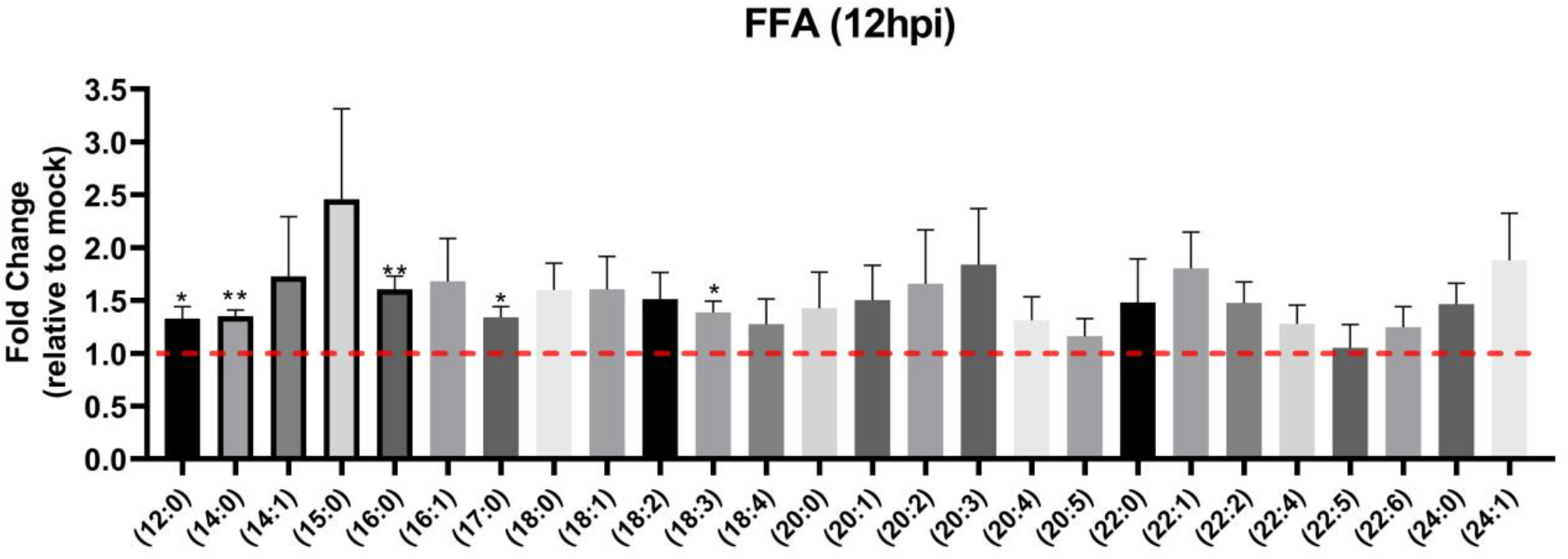
Quantitative lipidomics of saturated and unsaturated free fatty acids in mock vs MHV-68 infected cells. Mock (control) and MHV-68 infected NIH3T3 cells were harvested at 12 hpi (three independent experiments). All measured saturated and unsaturated free fatty acids (FAA) were graphed as relative fold change (y-axis) over mock using unpaired t-tests (p<0.05). Red dashed line at 1.0 indicates no change compared to mock infected cells.

**Table S1: Aqueous metabolomics normalized fold change time course in mock vs MHV-68 infected cells**.

**Table S2: Quantitative lipidomics normalized fold change time course in mock vs MHV-68 infected cells**.

